# Efficacy of dynamic eigenvalue in anticipating and distinguishing tipping points^†^

**DOI:** 10.1101/2024.01.09.574944

**Authors:** Kaustubh Kulkarni, Smita Deb, Partha Sharathi Dutta

## Abstract

The presence of tipping points in several natural systems necessitates having improved early warning indicators to provide accurate signals of an impending transition to a contrasting state while also detecting the type of transition. Various early warning signals (EWSs) have been devised to forecast the occurrence of tipping points, also called critical transitions. Dynamic eigenvalue (DEV) is a recently proposed EWS that can not only predict the occurrence of a transition but also certain types of accompanying bifurcations. Here, we study the effectiveness and limitations of DEV as an EWS for diverse kinds of critical phenomena. We demonstrate that DEV is a powerful EWS that shows promising results in anticipating catastrophic (first-order or discontinuous) and non-catastrophic (second-order or continuous) transitions in discrete and continuous dynamical systems. However, it falls short in the case of piecewise smooth systems. Further, the ability of DEV to forecast the type of transition is limited, as it cannot differentiate saddle-node bifurcation from transcritical and pitchfork bifurcations. Despite these limitations, we show that DEV can work as a robust indicator for varying rates at which the transition is approached and with systems involving colored noise.

## Introduction

Complex natural systems may undergo sudden abrupt changes in their state and conditions known as critical transitions (Carpenter and Brock 2006, Dakos et al 2008, Hirota et al 2011, Mumby et al 2007, Kéfi et al 2007). Critical transitions are often accompanied by local bifurcations and occur when the system crosses a critical threshold, i.e., a tipping point (Scheffer et al 2012). In the proximity of a tipping point, even a tiny perturbation to the system can lead to sudden and large changes in the system’s state. These transitions may shift the system to a new undesirable state from where a reversal might be practically impossible. Hence, it is crucial to detect such transitions in advance to undertake possible measures to prevent the shift or at least minimize its impact.

A stable equilibrium point of a dynamical system exhibits a positive rate of return in response to local perturbations. As the system approaches a bifurcation point where the equilibrium disappears or becomes unstable, this rate of return approaches zero (Carpenter et al 2008, Wissel 1984). This phenomenon is called critical slowing down (CSD) (van Nes and Scheffer 2007). CSD is the basis of most early warning signals (EWSs), which are statistical indicators that can anticipate forthcoming critical transitions. Numerous EWSs have been proposed in the literature, some of which include variance (Carpenter and Brock 2006), return rate (Carpenter et al 2008), autocorrelation at-lag-1 (Dakos et al 2008), and skewness (Guttal and Jayaprakash 2008). However, a transition may occur in the presence or absence of CSD (Hastings and Wysham 2010, Boettiger and Hastings 2013, Hillebrand et al 2020). It has been shown that these generic CSD-based indicators are more likely to precede catastrophic and non-catastrophic bifurcation-induced transitions that possess characteristics of CSD. They tend to be sensitive to the parameter values used while computing them, such as detrending bandwidth and sliding window size (Dutta et al 2018). Moreover, preexisting EWSs lack a quantitative estimation for detecting transitions, in the absence of which a signal cannot be undoubtedly called a true positive (Boettiger and Hastings 2012b). Thus, although these generic EWSs are supposed to be the precursors of massive and often calamitous transitions, they possess several limitations and may result in false signals (Boerlijst et al 2013, Dakos et al 2012a,b, Ditlevsen and Johnsen 2010, Dutta et al 2018, Hastings and Wysham 2010). It has been suggested that the knowledge of the mechanism behind a possible critical transition might be essential for utilizing EWSs (Dutta et al 2018). Yet, it remains a challenge to detect thresholds and forecast critical transitions from available empirical data when the underlying dynamical processes are unknown (Hillebrand et al 2020, O’Brien et al 2023).

Dynamic eigenvalue (DEV) is a newly introduced EWS that can potentially forewarn the occurrence of a transition as well as detect the underlying dynamical processes, i.e., the type of bifurcation involved (Grziwotz et al 2023). Mathematically, DEV is the dominant eigenvalue of the time-varying Jacobian matrix of the system estimated by deploying the S-map method (Deyle et al 2016, Sugihara et al 1994), an algorithm comprising lag-embedding empirical dynamic modeling (EDM) methods (Chang et al 2017, Deyle et al 2016, Ushio et al 2018, Ye et al 2015). Grziwotz et al (2023) demonstrates the applicability of DEV in predicting and classifying transitions using discrete dynamical models as well as empirical data. They have shown that as the system approaches the tipping point, |DEV| → 1, while the values approached by the real and the imaginary parts categorize the transition into three kinds, viz., saddle-node (fold) bifurcation (Re(DEV)→ 1, Im(DEV)→ 0), period-doubling bifurcation (Re(DEV)→ −1, Im(DEV)→ 0), and Neimark-Sacker bifurcation (|Re(DEV)| ↛ 1, Im(DEV) ↛ 0). This numerical threshold may provide DEV with an edge over the traditional EWSs, such as variance and autocorrelation at lag-1.

While Grziwotz et al (2023) have shown that DEV can forewarn and categorize certain types of catastrophic and non-catastrophic transitions, it is important to investigate the efficacy of DEV under diverse contexts to ascertain its utility as a reliable EWS. More importantly, it is necessary to establish how well it classifies the type of transition, which the traditional EWSs are unable to determine. Considering the myriad transitions that can occur in complex natural systems and the restricted amount of information the trend in an eigenvalue can provide, how effectively DEV anticipates the transitions and their type remains a significant question. Preexisting EWSs have some limitations (Boerlijst et al 2013, Dakos et al 2012a,b, Dutta et al 2018, Hastings and Wysham 2010), which become apparent when they are applied to real data that are often irregular and sparse. Hence, it is essential to explore the advantages and drawbacks of newly proposed EWSs before validating them as robust EWSs. Thus, we appraise the performance of DEV on a multifarious suite of discrete and continuous models. First, we use discrete models showing transcritical and pitchfork bifurcations to illustrate that they result in the same pattern in DEV as fold bifurcation, thus limiting the classification potential of DEV for fold bifurcation. On these models, we test the effect of the rate at which the transition is approached on DEV results since the “rate of forcing” is known to play its part (Clements and Ozgul 2016) in hampering the efficacy of early warning indicators. Further, we consider seven continuous dynamical models showing different kinds of transitions and probe the efficacy of DEV on these models. Additionally, we employ discrete models showing piecewise smooth bifurcations to explore whether this method can be generalized to piecewise smooth systems. This is particularly important as it is difficult to determine from empirical pre-transition data whether the associated dynamical processes pertain to smooth or piecewise smooth dynamical systems. Finally, we test whether DEV remains a robust indicator when the system entails colored (correlated) noise.

Overall, DEV is an effectual EWS that can presage the onset of several kinds of transitions, and its utility is not restricted to only a few types. Its numerical threshold makes it robust to the rate at which the transition is approaching, providing a valuable improvement over generic EWSs. The numerical threshold remains robust with correlated noise as well. However, as we show, it has a limited ability to distinguish the types of transitions from one another. Moreover, we observe that DEV cannot signal the occurrence of transitions when the associated dynamical process is piecewise smooth. In a nutshell, we critically assess the applicability of DEV as an EWS for different kinds of bifurcations in various systems. We emphasize that though DEV is an efficacious EWS correctly signaling the occurrence of different transitions, one should conclude about the ensuing dynamics only after a careful interpretation of its results. We underline the importance of taking into account multiple different EWSs to forecast critical transitions, as each EWS comes with its advantages and disadvantages.

## Models and Methods

### DEV calculations and analysis

DEV is largely based on the bifurcation theory and derived using the empirical dynamic modeling (EDM) framework (Deyle et al 2016, Grziwotz et al 2023, Sugihara et al 1994, Ushio et al 2018). EDM aids in rendering an impression of the attractor of the system using lags from the time series of a single variable. This reconstructed attractor retains the fundamental properties of the system (Deyle et al 2016, Ushio et al 2018, Ye et al 2015). DEV uses an EDM method known as the S-map (Deyle et al 2016, Sugihara et al 1994) to estimate the time-varying Jacobian of the system.

The algorithm to calculate DEV is as follows: We begin with a time series {*x*_1_, *x*_2_, …, *x*_*T*_ }. We divide the time series into sliding windows of size *w*, which slide with a step size *s*. Within each window, we construct a univariate lag embedding with an embedding dimension *E* and time lag *τ* . The parameters *w, s, E*, and *τ*, are to be chosen separately. Simply put, we write the variable values at time point *X*_*t*+*τ*_ = [*x*_*t*+*τ*_, *x*_*t*_, *x*_*t*−*τ*_, …, *x*_*t*−(*E*−2)*τ*_ ]^*T*^ as a function of the values at *X*_*t*_ = [*x*_*t*_, *x*_*t*+*τ*_, …, *x*_*t*−(*E*−2)*τ*_, *x*_*t*−(*E*−1)*τ*_ ]^*T*^, i.e., *X*_*t*+*τ*_ = *g*(*X*_*t*_), where [… ]^*T*^ stands for transpose of the row vector. For this reconstructed system, we estimate the time-varying local Jacobians, **Ĵ**(*t*), such that *X*_*t*+*τ*_ = **Ĵ***X*_*t*_ + *C*. The dynamics of this reconstructed system are thus represented as follows:

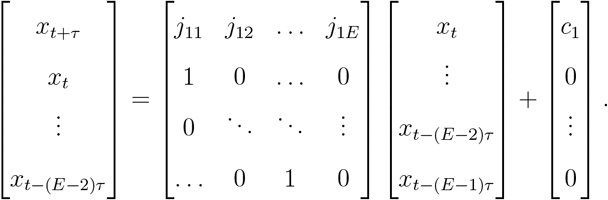

The time-varying Jacobian matrices are estimated from the S-map algorithm (Deyle et al 2016, Sugihara et al 1994). The dominant eigenvalue of this Jacobian matrix is called the dynamic eigenvalue (DEV) (Grziwotz et al 2023). The S-map method essentially performs locally weighted linear regression to obtain the coefficients *j*_11_, *j*_12_, …, *j*_1*E*_ at a given time point (Deyle et al 2016). The term *c*_1_ in the vector *C* is the constant term obtained in weighted linear regression.

As the system approaches the bifurcation point, |DEV| approaches 1. Note that even for continuous systems, |DEV| will approach 1 since the data obtained after numerical integration are discrete. Importantly, the DEV is not an eigenvalue of the actual dynamical system. It is the dominant eigenvalue of the Jacobian of a discrete imprint of the system constructed using lag embedding.

Similar to the univariate DEV approach explained above, we can also calculate DEV using multivariate S-maps (Deyle et al 2016, Grziwotz et al 2023). In that case, the embedding dimension *E* corresponds to the number of variables in the time series. For example, for two variables *x* and *y*, we would write the system as:

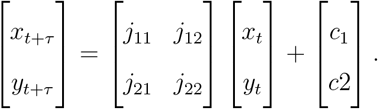

From our preliminary investigation of the factors affecting DEV trends, we observed that the window size *w* impacts the smoothness of the trends obtained. A higher window size results in smoother changes in DEV but is likely to overfit the data (Grziwotz et al 2023). Thus, in every case, we chose an appropriate window size that resulted in reasonably smooth trends. We fixed the step size *s* = *w/*5 throughout the analysis. We first chose *E* and *τ* by trial-and-error after observing the DEV results and then performed sensitivity analysis (see *Sensitivity analysis*) on *E* and *τ* in discrete models with white noise (*SI Appendix*, Table S1) to illustrate the influence of these two parameters on the results obtained.

### Discrete dynamical systems

Here, we examined three discrete models showing fold, transcritical, and pitchfork bifurcations, viz., the Noy-Meir Model (Noy-Meir 1975), the Lotka-Volterra Model (Neubert and Kot 1992) and the reduced Lorenz Model (Elabbasy et al 2014, Lorenz 1989, Whitehead and MacDonald 1984), respectively. The rationale behind considering these bifurcations is that for all of them, the dominant eigenvalue is real and crosses the DEV threshold of 1 as the bifurcation occurs. We aimed to infer whether DEV can distinguish between these three types of bifurcations, out of which fold bifurcation is catastrophic. In contrast, the remaining two can be grouped as being non-catastrophic. The model equations and parameter values are given in Table S1 (see *SI Appendix*). For each of these models, we considered Gaussian white noise, such that the models took the form:

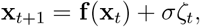

where *ζ*_*t*_ ∼ *N* (0, 1) is drawn from the standard normal distribution and *σ* is the amplitude of the noise.

### Continuous dynamical systems

We analyzed seven continuous-time ecological models (Dutta et al 2018), listed in Table S2 (*SI Appendix*) along with the parameter values. The first of them shows a fold (saddle-node) bifurcation (Ludwig et al 1978, May 1977, Noy-Meir 1975) (Table S2, Model 1), whereas with another set of parameters, it shows no sharp transition but a gradual change (Table S2, Model 6). It is essential to consider a model that does not show a transition because focusing only on systems undergoing transitions can underestimate the rate of false positives (Boettiger and Hastings 2012a,b). For a third set of parameters, this model exhibits a sharp transition without a bifurcation (Table S2, Model 7). Two modified versions of the aforementioned model show transcritical (Table S2, Model 2) and pitchfork (Table S2, Model 4) bifurcations, respectively. We also consider a model exhibiting supercritical Hopf bifurcation (Rosenzweig 1971) (Table S2, Model 3) and a model that undergoes bifurcation without CSD (Schreiber and Rudolf 2008) (Table S2, Model 5). We incorporated Gaussian white noise into each of these models. Thus, these continuous time stochastic models take the following general form:

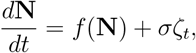

where *f* (**N**) is the deterministic skeleton and *ζ*_*t*_ ∼ *N* (0, 1) is drawn from the standard normal distribution. The parameter *σ* determines the amplitude of the noise. For simplicity in Model 5, we added noise only to one variable as different variables had different orders of magnitude.

### Piecewise smooth discrete dynamical systems

We also examined a discrete model (a map) that shows various piecewise smooth transitions under different parameter settings, viz., no bifurcation, fold bifurcation, period-doubling bifurcation, and transition from a period-1 orbit to chaos (Banerjee et al 2000). We also considered a discrete model with a piecewise smooth Neimark-Sacker bifurcation (De et al 2012). The Neimark-Sacker bifurcation is the discrete system equivalent of the Hopf bifurcation. Table S3 (*SI Appendix*) lists the equations and parameter values used. In the aforesaid piecewise smooth models as well, we incorporated Gaussian white noise as in the smooth discrete examples (see *Discrete dynamical systems*).

### Discrete dynamical systems with colored noise

We further aimed to investigate the effect of noise correlation on DEV results. For this purpose, we considered the same three models examined used by Grziwotz et al (2023), but incorporated colored noise (see *SI Appendix*, Eqn. (S1)) in them. These models, viz., the Noy-Meir Model, the Hénon Map, and the Rosenzweig-MacArthur Model, show fold, period-doubling, and Neimark-Sacker bifurcations, respectively. The model equations and parameter values are given in Table S4 (*SI Appendix*). In each of these three models, we added colored noise to the variable for which DEV was evaluated, as described in Section S3 (*SI Appendix*, S3: Models).

For the numerical simulation of each of the models in each scenario, we obtained a time series of length 10, 000 by incrementing the control parameter with time. The continuous models were numerically integrated with an integration time step of *dt* = 0.01. We evaluated DEV from the simulated time series. DEV parameters are given in Tables S5, S6, S7, and S8 (see *SI Appendix*). On the smooth discrete models (*SI Appendix*, Table S1), we also tested how the rate of forcing affects the results given by DEV. For this purpose, we generated time series by incrementing (decrementing in the case of Model 2 in Table S1) the control parameter from its initial value to 10 different values (Clements and Ozgul 2016). For each of these 10 values, we generated 100 time series and performed DEV analysis on each of them. Note that the pre-transition time series had different lengths for different final values of the control parameter. All the simulations and computations were performed in `R (version 4.3.2)` (R Core Team 2022). S-maps were implemented using the `rEDM` library `(version 1.14.0)` (Park et al 2023).

## Results

### DEV trends for catastrophic and non-catastrophic transitions

DEV calculated for pre-transition data generated from the considered discrete models of fold, transcritical, and pitchfork bifurcations shows an increasing trend signaling an upcoming transition in each of the cases (Fig. 1A). However, in all these cases, the characteristics of DEV remain the same, i.e., |DEV| → 1, Re(DEV)→ 1, and Im(DEV)→ 0 (Figs. 1B-1D). Thus, DEV cannot distinguish between these three kinds of transitions. Hence, Grziwotz et al (2023)’s assertion that DEV can tell apart fold bifurcation, period-doubling bifurcation, and Neimark-Sacker bifurcation from each other could be appropriately rephrased as follows: Though the latter two kinds of bifurcations (period-doubling and Neimark-Sacker) are successfully distinguished, a DEV trend noted by Grziwotz et al (2023) for fold bifurcation could be a warning of one of at least three different types, viz., fold bifurcation, transcritical bifurcation, and pitchfork bifurcation. This is important in light of the knowledge that fold bifurcation is a catastrophic bifurcation (resulting in a discontinuous transition), whereas transcritical and pitchfork bifurcations belong to the non-catastrophic class (resulting in a continuous transition) (Kéfi et al 2013, Dutta et al 2018). Thus, a system showing the DEV output mentioned by Grziwotz et al (2023) for fold bifurcation need not necessarily be nearing a catastrophic transition but could be advancing towards a non-catastrophic reversible transition as well. In Fig. 1 (and similarly in Figs. 3 and 4), the DEV trends have been averaged over 100 different realizations of simulated time series generated from the considered models each, as DEV from a single time series shows more variability in the trends obtained.

**Figure 1.**
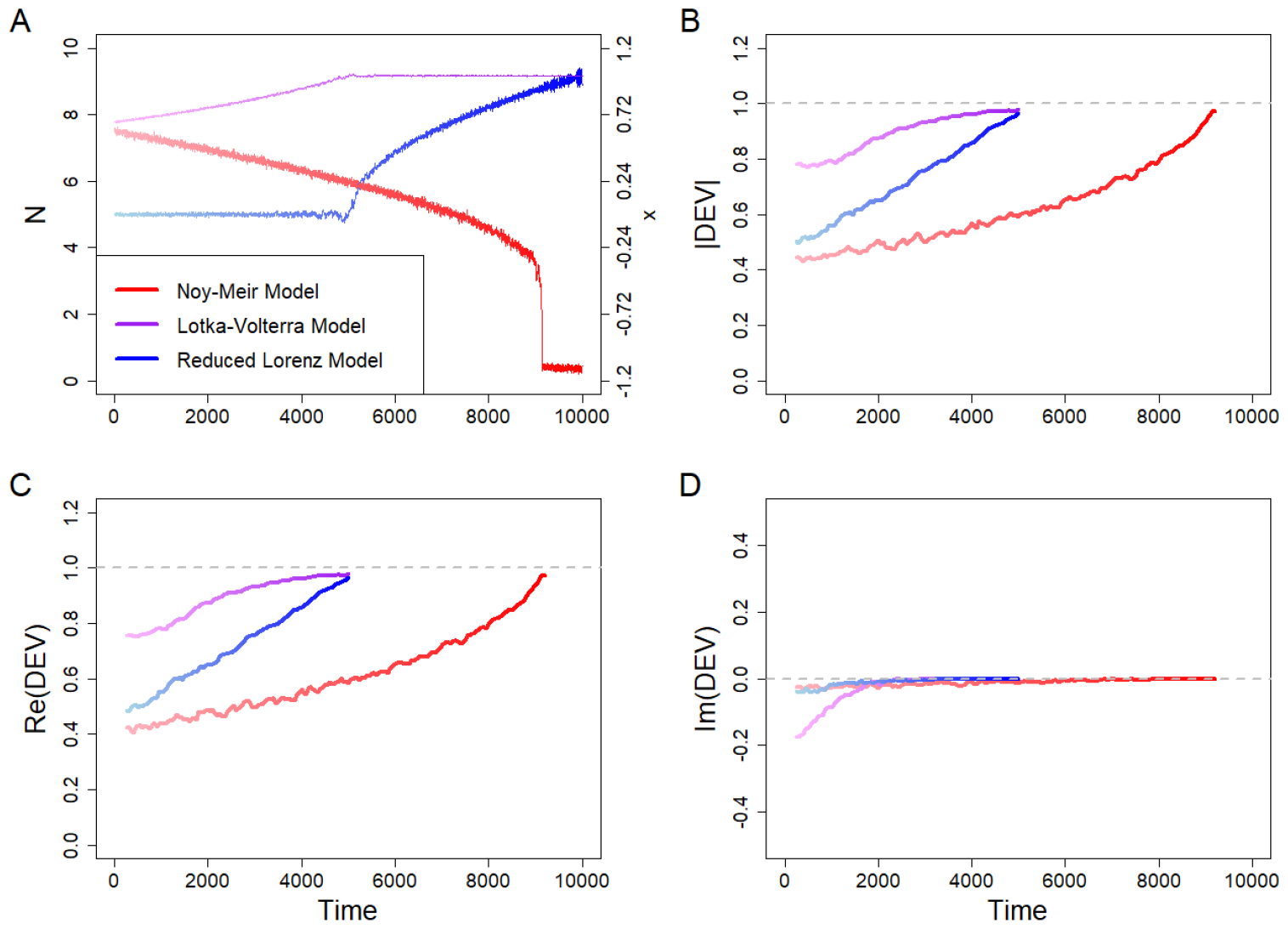
(A) Time series, (B) |DEV|, (C) Re(DEV), and (D) Im(DEV) for the Noy-Meir Model (fold bifurcation), the Lotka-Volterra Model (transcritical bifurcation), and the reduced Lorenz Model (pitchfork bifurcation) in discrete dynamical systems (see *SI Appendix*, Table S1). The time series is a single realization, whereas the DEV values averaged over 100 realizations. The DEV parameter values used are given in *SI Appendix*, Table S5. All three models show similar DEV trends before the bifurcation, precluding us from distinguishing between them.

### Efficacy of DEV with fast system dynamics

The rate at which the transition approaches in a system, i.e., the rate of forcing, has been previously shown to influence the strength of generic EWSs (Clements and Ozgul 2016, van der Bolt et al 2021). In Figs. 2A, 2C, and 2E, we depict violin plots with |DEV| at the transition point for fold, transcritical, and pitchfork bifurcations. |DEV| reaches values close to 1 for all rates of forcing for all three models. Hence, the numerical threshold of DEV is unaffected by the rate at which a transition approaches. The second column (Figs. 2B, 2D, and 2F) displays the difference between the |DEV| value at the transition point and that at the beginning of the time series, denoted by Δ|DEV|. Δ|DEV| tells us the effectiveness of DEV as a qualitative EWS (Grziwotz et al 2023). For the fold bifurcation, Δ|DEV| has decreased with increasing rate of forcing, i.e., for high rates of forcing, Δ|DEV| was close to 1 even at the beginning of the time series. However, this could, in part, be caused by keeping the window size constant across the time series for different rates, where the initial trends get averaged out for shorter time series (i.e., for higher rates of forcing). Nevertheless, this averaging may also inform us about the faster rate of approach of the transition. Δ|DEV| is not remarkably influenced by the rate of forcing in transcritical and pitchfork bifurcations (Figs. 2D and 2F). Altogether, DEV appears to be robust to the rate of forcing for fold, transcritical, and pitchfork bifurcations.

**Figure 2.**
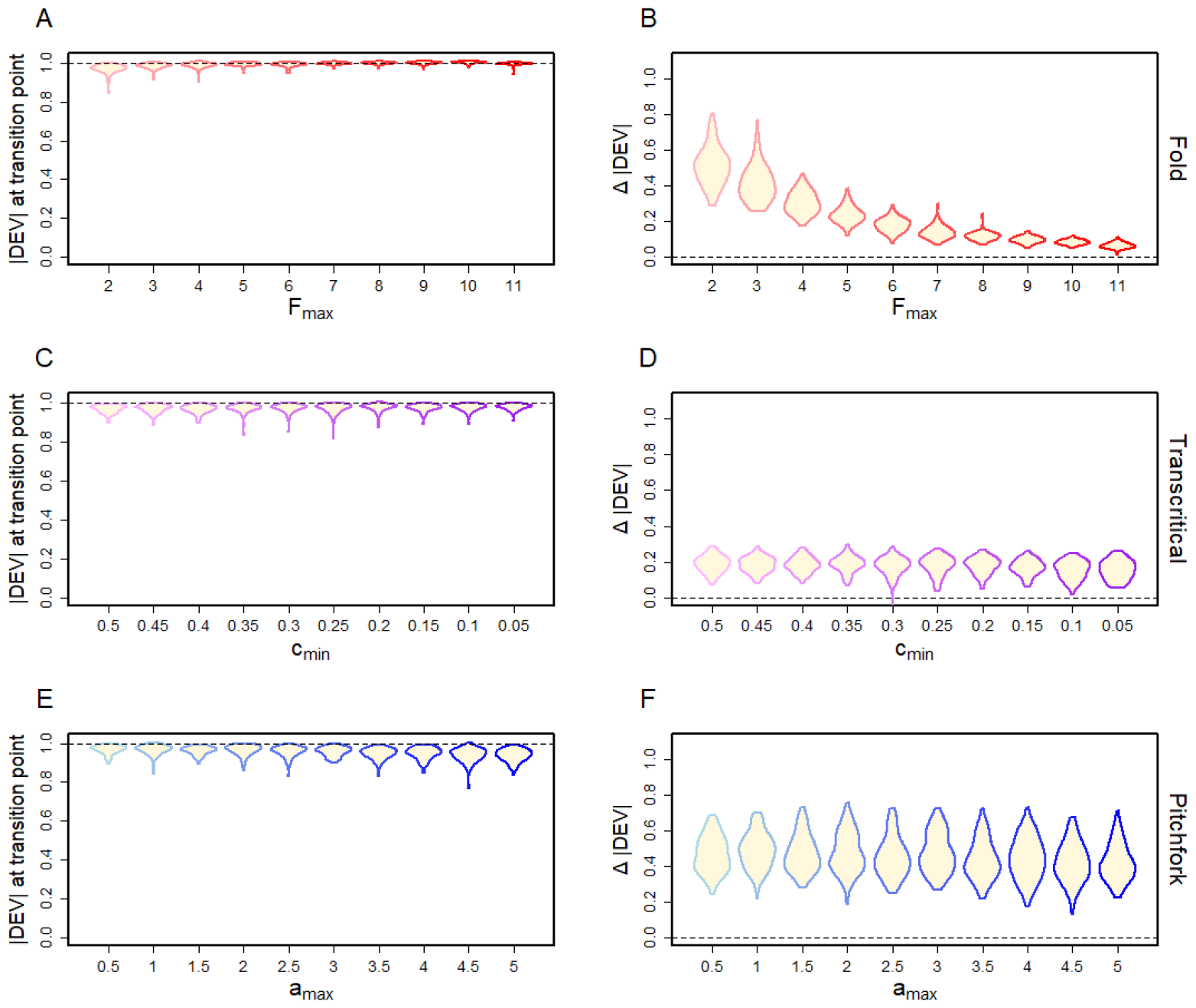
Violin plots showing the effect of rate of forcing on DEV results with discrete models (*SI Appendix*, Table S1) exhibiting: (A, B) Fold bifurcation, (C, D) Transcritical bifurcation, and (E, F) Pitchfork bifurcation. The rate of forcing increases from left to right in every plot. The first column shows |DEV| values at the transition point, and the second column shows Δ |DEV| (the difference between |DEV| at the transition point and that at the beginning of the time series). Each violin plot is based on 100 simulated time series. The DEV parameter values used are given in *SI Appendix*, Table S5. The |DEV| threshold is robust to the rate at which the transition approaches, and so is Δ |DEV|, except in the Noy-Meir Model (fold bifurcation), where it decreases with increasing rate of forcing.

### DEV for continuous dynamical systems exhibiting a variety of transitions

The continuous models we considered (see *SI Appendix*, Table S2) show various transitions (*see Methods, Continuous dynamical systems*). DEV signals some of these transitions, with |DEV| reaching the numerical threshold of 1 at the tipping points. Fig. 3 shows the time series, absolute values, and real and imaginary parts of DEV for all the cases. Case-by-case explanations are given in Table 1.

**Figure 3.**
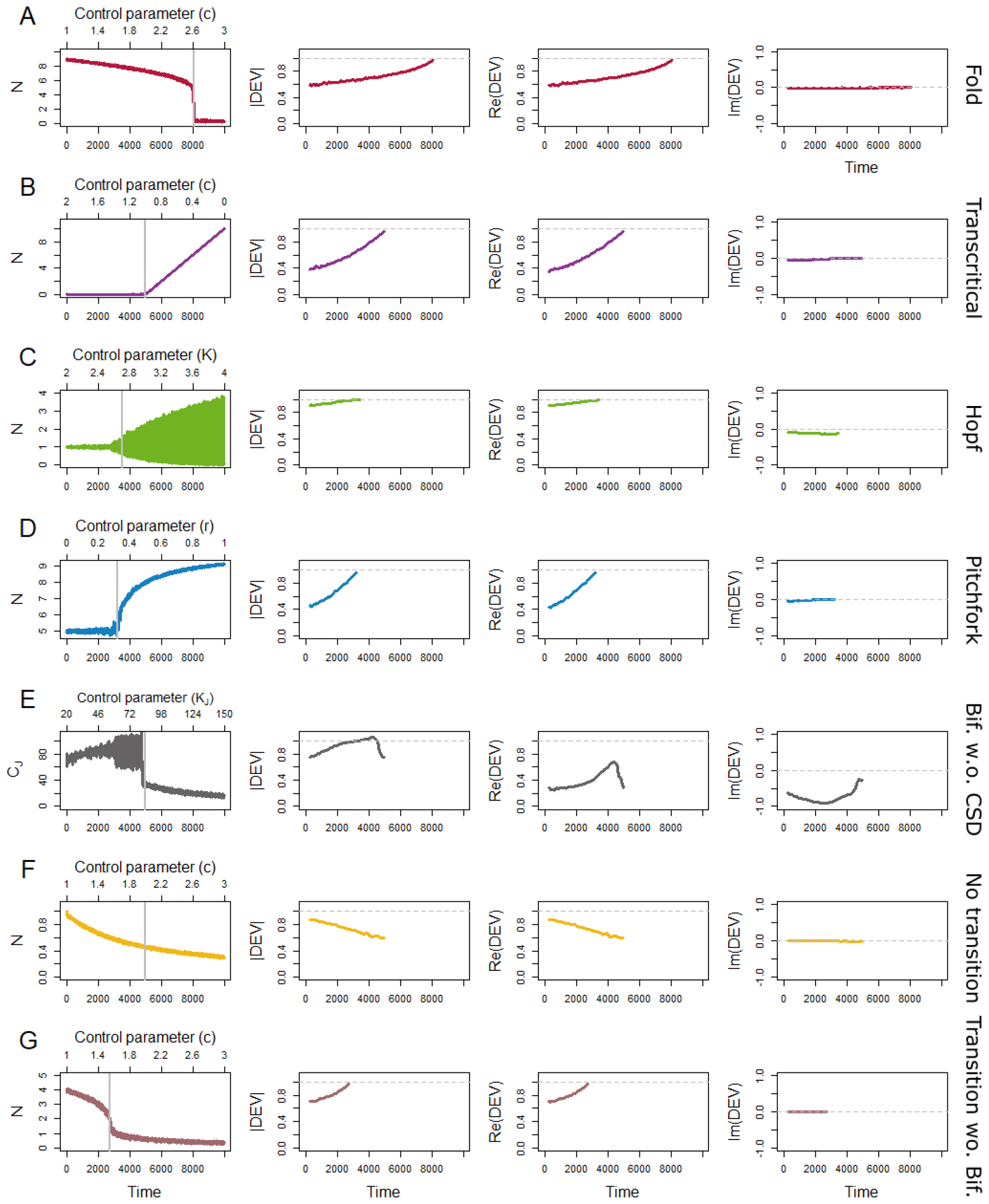
Time series, |DEV|, Re(DEV), and Im(DEV) for: (A) fold bifurcation, (B) transcritical bifurcation, (C) supercritical Hopf bifurcation, (D) pitchfork bifurcation, (E) bifurcation without CSD, (F) no transition, and (G) transition without bifurcation in continuous dynamical systems (for models, see *SI Appendix*, Table S2). The time series is a single realization, whereas the DEV values averaged over 100 realizations. Grey vertical lines in each sub-figure in the left column mark proximity to the transition points. The DEV parameter values used are given in *SI Appendix*, Table S6. Bifurcations/transitions in sub-figures A, B, D, and G cannot be distinguished from one another, while the Hopf bifurcation in sub-figure C can be separated from the rest. Transition without a bifurcation (E) is not signaled, and so is the case where there is no transition (F).

**Table 1.** DEV results for continuous models showing various types of transitions: (1) Fold bifurcation, (2) transcritical bifurcation, (3) pitchfork bifurcation, and (7) the transition without a bifurcation are all predicted by DEV. These transitions show similar DEV trends and thus cannot be distinguished from one another. (3) Supercritical Hopf bifurcation is successfully anticipated and can be distinguished from the remaining types. (5) In the model showing a bifurcation without CSD, the transition is not predicted. (6) In the case of the model with no transition, a transition is not predicted (see *SI Appendix*, Table S2).

| Models | Type of system | DEV Results | Increasing trend signaling transition | Indistinguishable from models | Fig. |
| --- | --- | --- | --- | --- | --- |
| 1. | Fold bifurcation | $\vert \text{DEV}\vert \rightarrow 1$<br>$\text{Re}(\text{DEV}) \rightarrow 1$<br>$\text{Im}(\text{DEV}) \rightarrow 0$ | Yes | 2, 4, 7 | **3A** |
| 2. | Transcritical bifurcation | $\vert \text{DEV}\vert \rightarrow 1$<br>$\text{Re}(\text{DEV}) \rightarrow 1$<br>$\text{Im}(\text{DEV}) \rightarrow 0$ | Yes | 1, 4, 7 | **3B** |
| 3. | Supercritical Hopf bifurcation | $\vert \text{DEV}\vert \rightarrow 1$<br>$\text{Re}(\text{DEV}) \not\rightarrow 1$<br>$\text{Im}(\text{DEV}) \not\rightarrow 0$ | Yes | None | **3C, S1** |
| 4. | Pitchfork bifurcation | $\vert \text{DEV}\vert \rightarrow 1$<br>$\text{Re}(\text{DEV}) \rightarrow 1$<br>$\text{Im}(\text{DEV}) \rightarrow 0$ | Yes | 1, 2, 7 | **3D** |
| 5. | Bifurcation without CSD | $\vert \text{DEV}\vert \not\rightarrow 1$<br>$\text{Re}(\text{DEV}) \not\rightarrow 1$<br>$\text{Im}(\text{DEV}) \neq 0$ | No | NA | **3E, S2** |
| 6. | No transition | $\vert \text{DEV}\vert \not\rightarrow 1$<br>$\text{Im}(\text{DEV}) = 0$ | No (as desired) | NA | **3F** |
| 7. | Sharp transition without bifurcation | $\vert \text{DEV}\vert \rightarrow 1$<br>$\text{Re}(\text{DEV}) \rightarrow 1$<br>$\text{Im}(\text{DEV}) \rightarrow 0$ | Yes | 1, 2, 4 | **3G** |

Our results illustrate that DEV is a useful EWS that can forewarn the occurrence of transitions not only in discrete systems (Grziwotz et al 2023) but also for a large variety of transition types in continuous systems. However, at the same time, it is evident from the above analysis that DEV shows similar results for fold bifurcation (Fig. 3A), transcritical bifurcation (Fig. 3B), pitchfork bifurcation (Fig. 3D), and transition without a bifurcation (Fig. 3G) (see Table 1). Thus, DEV is unable to distinguish between these four instances of transitions, though it can distinguish Hopf bifurcation from the remaining types. This drawback is rooted in the bifurcation theory, wherein the properties shown by eigenvalue during these transitions are similar (Bury et al 2020).

The results for Hopf bifurcation are similar for both the variables in the system (Fig. 3 C, *SI Appendix*, Fig. S1A) and multivariate DEV also shows a positive signal (*SI Appendix*, Fig. S1B). In the model without a transition, |DEV| does not approach 1 and hence, signaling no transition (Fig. 3F), as is desirable. The inability of DEV to detect the bifurcation, not characterized by CSD (Fig. 3E), is a limitation of the method. Here, an increasing trend is observed, but |DEV| crosses 1 before the transition. It again decreases and falls below 1 prior to the transition. Im(DEV) is non-zero. None of the other variables in the system are able to signal the upcoming transition (*SI Appendix*, Fig. S2A-S2C), and nor does multivariate DEV do so (*SI Appendix*, Fig. S2D). Im(DEV) shows a spike to zero in some cases (*SI Appendix*, Figs. S2A and S2D).

### DEV for piecewise smooth systems

DEV cannot anticipate transitions occurring in piecewise smooth dynamical systems. In these models, the functions describing the system abruptly change after the transition point (*see SI Appendix, Table S3*). |DEV| does not show any trend in such cases, but shows a spike at the bifurcation point. Hence, right after the transition, we might be able to determine the type of transition the system has undergone, based on the real and imaginary parts of DEV. If the Im(DEV) is non-zero, it’s likely to be either a Neimark-Sacker bifurcation or a transition to chaos. If Im(DEV) is zero, and Re(DEV) is −1, the system is likely to have undergone a period-doubling bifurcation. DEV may thus be able to work as a post-transition classification method in these cases, though not as EWS. Case-by-case explanations are given in Table 2.

**Table 2.** DEV results for piecewise smooth models showing: (1) No bifurcation, (2) fold bifurcation, (3) period-doubling bifurcation, (4) transition from a Period-1 orbit to chaotic dynamics, and (5) Neimark-Sacker bifurcation (see *SI Appendix*, Table S3). The transitions in these models occur abruptly, as the functional form of the dynamics changes as the tipping point is crossed.

| Models | Type of system | DEV Results | Increasing trend signaling transition | Fig. |
| --- | --- | --- | --- | --- |
| 1. | No bifurcation | No trend, sharp transition at critical point | No | 4A |
| 2. | Fold bifurcation | No trend, $\text{Im}(\text{DEV})$ is 0 | No | 4B |
| 3. | Period doubling bifurcation | No trend, $\vert \text{DEV}\vert $ spikes to a value $> 1$ at transition point; $\text{Re}(\text{DEV})$ spikes to less than -1 | No | 4C |
| 4. | Period-1 to Chaos | No trend, $\vert \text{DEV}\vert $ spikes to $> 1$, $\text{Re}(\text{DEV})$ spikes to $-1$, $\text{Im}(\text{DEV})$ spikes to a negative value | No | 4D |
| 5. | Neimark-Sacker bifurcation | $\vert \text{DEV}\vert $ shows no trend, dips at the transition point, and rises again; $\text{Im}(\text{DEV})$ dips from 0 to a negative value | No | 4D, S3 |

Thus, in piecewise smooth discrete systems, calculating DEV by perturbating the bifurcation parameter is unable to provide any indication of the occurrence of the bifurcation. To anticipate bifurcation-induced transitions in a piecewise smooth system, one possible way might be to perform regularization to incorporate smoothness assumptions in the present framework of DEV computation (Makarenkov and Lamb 2012).

### DEV for discrete systems with colored noise

We considered colored noise in the three models (Grziwotz et al 2023) undergoing fold, period-doubling, and Neimark-Sacker bifurcations by adding positively correlated noise to one variable in each model (see *SI Appendix*, Section S3: Models, Eqn. (S1)). The |DEV| threshold of 1 is robust to noise correlation in these models (see Figs. 5A, 5C, and 5E).

**Figure 4.**
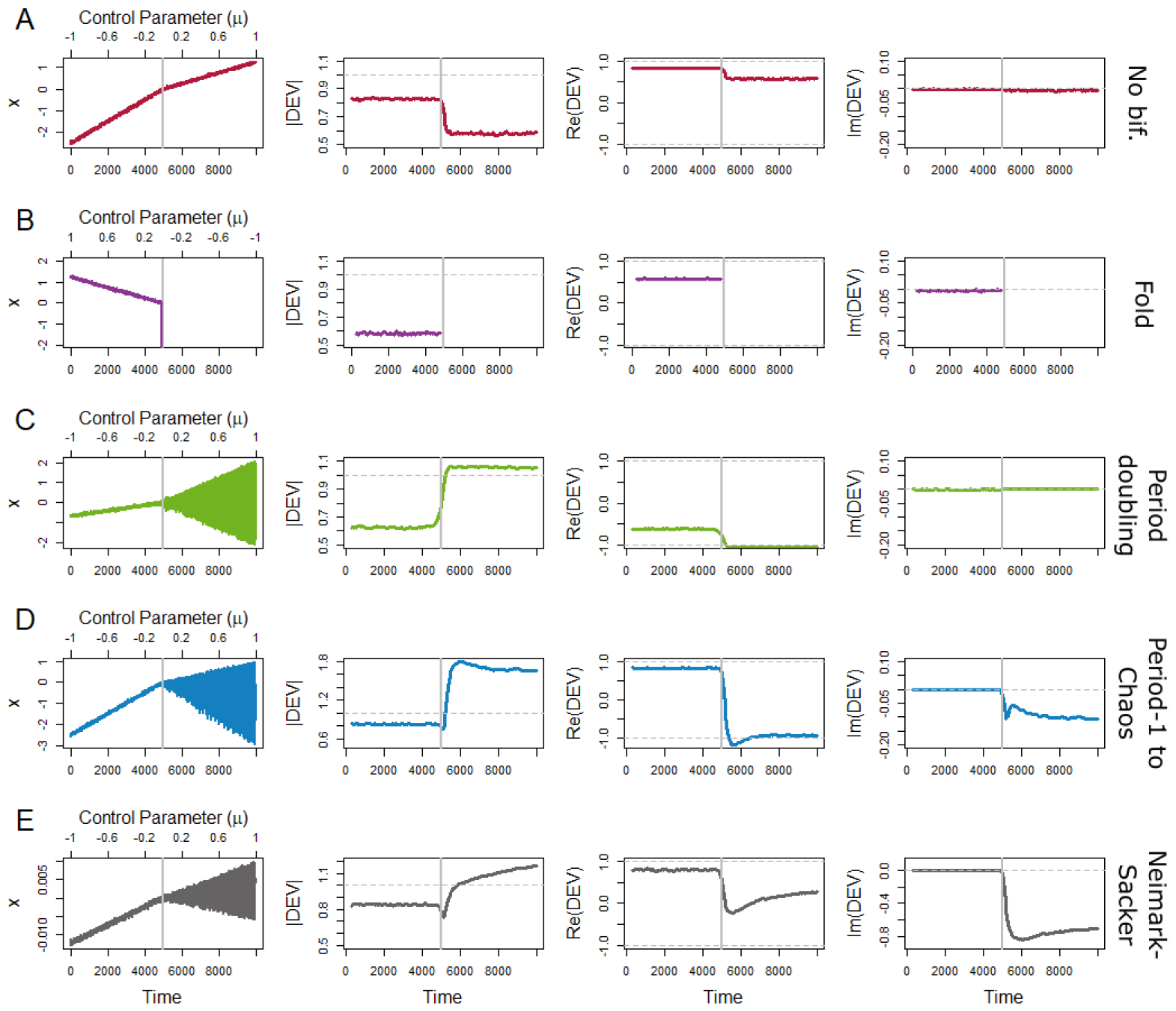
Time series, |DEV|, Re(DEV) and Im(DEV) for (A) No bifurcation, (B) Fold bifurcation, (C) Period doubling bifurcation, (D) Period-1 to chaos and (E) Neimark-Sacker bifurcation in piecewise smooth discrete dynamical systems (*SI Appendix*, Table S3). The time series is a single realization, whereas the DEV values averaged over 100 realizations. Grey vertical lines mark the transition points. The DEV parameter values used are given in *SI Appendix*, Table S7. None of the transitions is successfully predicted by DEV.

**Figure 5.**
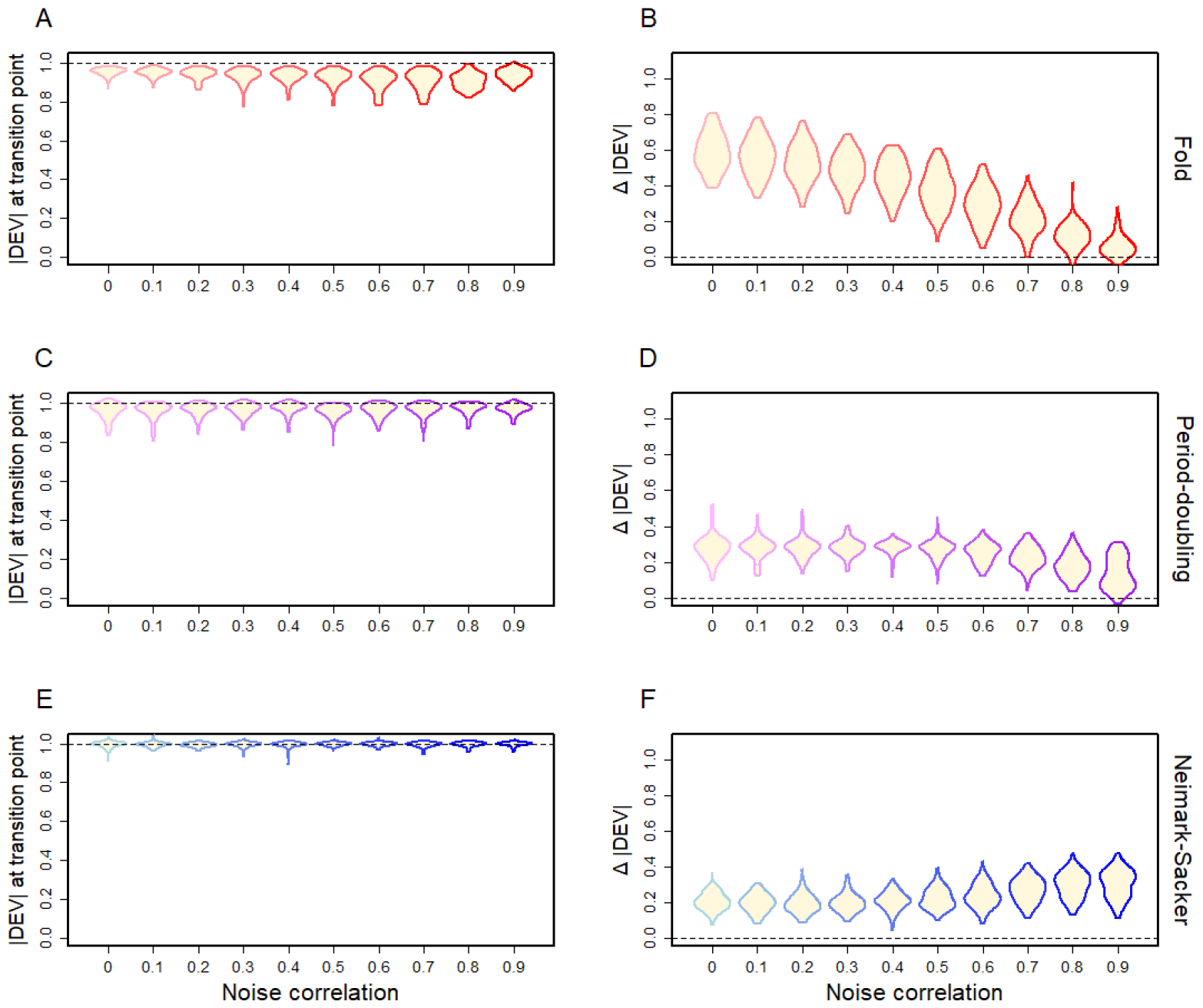
Violin plots showing the effect of noise correlation on DEV results for discrete models (*SI Appendix*, Table S4) with colored noise exhibiting: (A, B) Fold bifurcation, (C, D) Period-doubling bifurcation, and (E, F) Neimark-Sacker bifurcation. The first column (A, C, E) shows |DEV| values at the transition point, and the second column (B, D, F) shows Δ |DEV| (the difference between |DEV| at the transition point and at the beginning of the time series). Each violin plot is based on 100 simulated time series. The DEV parameter values are given in *SI Appendix*, Table S8. The DEV threshold is robust to noise correlation values, and so is Δ |DEV|, except in the case of the Noy-Meir Model (fold bifurcation), where it decreases with increasing noise correlation.

However, Δ|DEV|, the difference between the |DEV| value at the transition point and at the beginning of the time series varied with noise correlation (Figs. 5B, 5D, and 5F). For fold bifurcation, Δ|DEV| decreased with increasing noise correlation, with higher noise correlation producing even negative Δ|DEV| values (Fig. 5B). For period-doubling bifurcation, while the distribution of Δ|DEV| also moved towards lower values with increased noise correlation (Fig. 5D), the decrease was less pronounced compared to fold bifurcation. Contrarily, Δ|DEV| was robust to noise correlation in the case of Neimark-Sacker bifurcation, even showing an increasing trend as noise correlation was increased (Fig. 5F). Thus, these results point out that colored or correlated noise can undermine the ability of DEV to show a trend, at least in some cases, while the numerical threshold is likely to remain robust concerning noise correlation.

### Sensitivity analysis

To observe how our chosen DEV parameters *E* and *τ* affect the results in the discrete models with white noise (*SI Appendix*, Table S1), we simulated 100 time-series for each model and performed DEV analysis for different values of *E* and *τ* (1 to 10). Then, we averaged the 100 DEV trends for each *E* − *τ* combination. The parameters we chose were among the best combinations w.r.t. the DEV threshold of 1 and Δ|DEV|. Fig. 6 and Figs. S4-S5 (*SI Appendix*) show heat-maps with |DEV| at transition point and Δ|DEV| for different values of *E* and *τ* . The degree of robustness to parameters varied across models.

**Figure 6.**
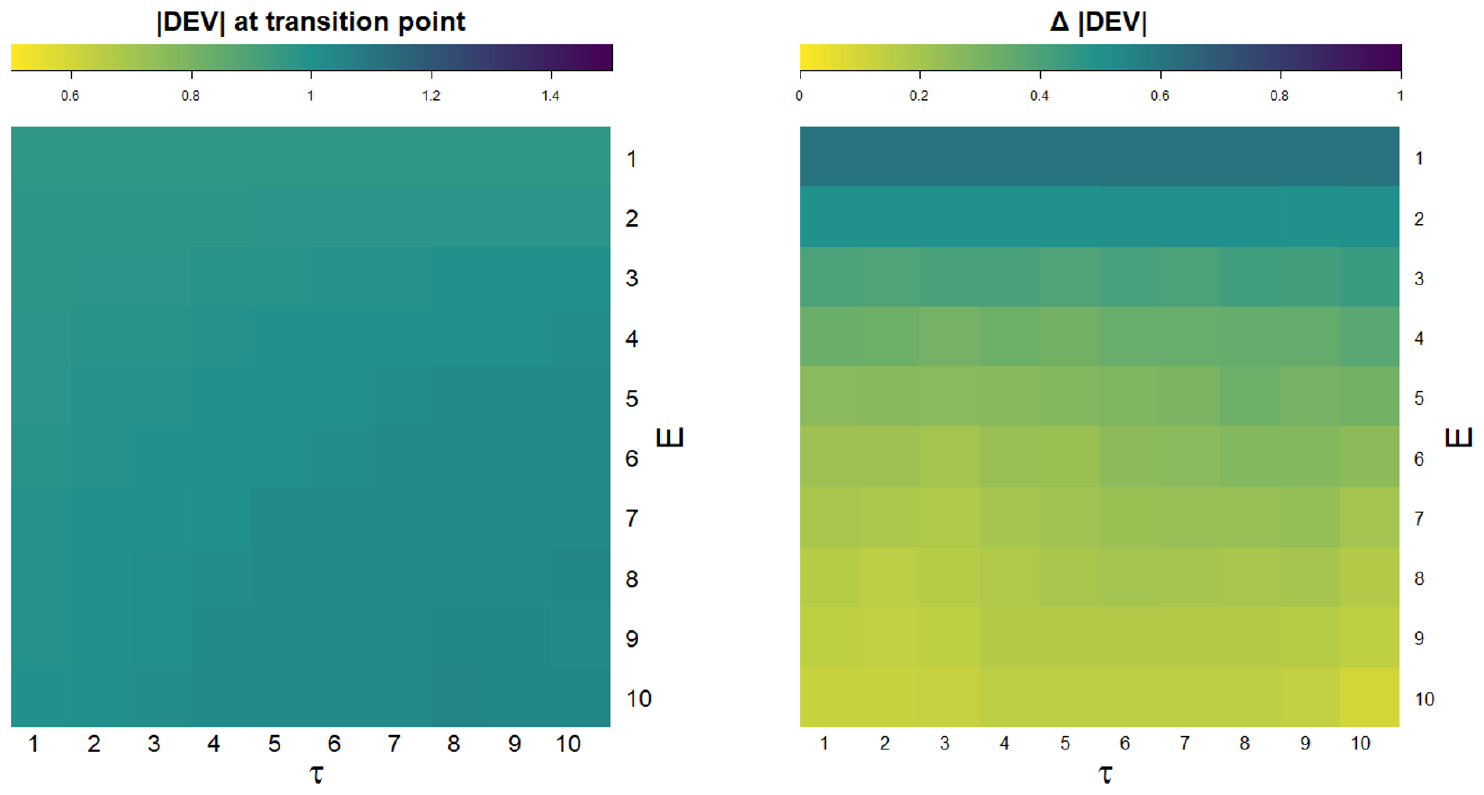
|DEV| at the transition point and Δ |DEV| for different combinations of *E* and *τ* for the discrete Noy-Meir Model (fold bifurcation) (see *SI Appendix*, Table S1). In the left figure, as the color changes from yellow to green to violet, |DEV| at the transition point increases from 0.5 to 1 to 1.5. |DEV| of 1 at the transition point is ideal. In the right figure, for the same color changes, Δ |DEV| increases from 0 to 0.5 to 1. Higher Δ |DEV| implies a qualitatively better signal. |DEV| at the transition point varies slightly with chosen parameters in the case of fold bifurcation.Δ |DEV| decreases with increasing *E*, and shows an increasing pattern with increasing *τ* .

The |DEV| at the transition point in the Noy-Meir Model (fold bifurcation) (Fig. 6) and the reduced Lorenz Model (pitchfork bifurcation) (*SI Appendix*, Fig. S5) showed minor sensitivity to the choice of *E* and *τ* . However, in both models, Δ|DEV| showed a degradation where it decreased with increasing values of *E*, and was more sensitive to *E* than *τ* . In the reduced Lorenz Model, it increased with increasing values of *τ* as well (*SI Appendix*, Fig. S5). In the Lotka-Volterra Model (transcritical bifurcation), |DEV| and Δ|DEV| (*SI Appendix*, Fig. S4) were more sensitive to the parameter choice than the other two models. The results showed that DEV trends became less and less effective with increasing values of both *E* and *τ*, with almost no trend being observed for high *E* and *τ* .

## Discussion

Dynamic eigenvalue (DEV) is a promising tool for its ability to anticipate critical transitions using a quantitative threshold and categorize an impending transition in smooth dynamical systems. We have shown that this ability of DEV is not limited to data obtained from discrete systems but is extendable to continuous systems. At the same time, we have also shown that DEV can classify bifurcations only up to a certain extent and cannot differentiate catastrophic transitions from non-catastrophic ones. This is vital because the urgency, the extent, and the manner of preventive action to be taken may depend on whether the approaching transition is catastrophic or non-catastrophic (Villa Martín et al 2015, Deb et al 2022).

The inability of DEV to discriminate among fold, transcritical, and pitchfork bifurcations stems from the bifurcation theory, wherein the analytical dominant eigenvalue shows the same trends for these three bifurcations. In all these three cases, the magnitude of the analytical dominant eigenvalue reaches 1, with the real part reaching 1 and the imaginary part equal to zero. The same is reflected in the trends in DEV. As Grziwotz et al (2023) show, DEV estimates the dominant eigenvalue of the Jacobian matrix of the system. DEV resembles the analytical eigenvalue when the S-map skill is high. Thus, as the analytical dominant eigenvalue shows the same trends for the aforementioned three types of bifurcations, DEV, too, follows the same lines, preventing a straightforward interpretation of its output.

As previously reported, generic EWSs fail to be consistent across different transition rates (Clements and Ozgul 2016, van der Bolt et al 2021); however, DEV is robust to the rate at which a system advances towards a transition. This is primarily because we can interpret DEV quantitatively, unlike other EWSs. Though higher rates of forcing may undermine the increasing trends shown by DEV, they have little effect on DEV at the tipping point. This is another advantage of DEV over generic EWSs, conferred upon by its numerical threshold. Similarly, the DEV threshold is robust to noise correlation; however, high noise correlation can weaken the increasing trends in |DEV|.

Though the concept of DEV is based on the eigenvalue patterns in discrete systems, we have demonstrated that DEV can be applied even to continuous systems. Though in continuous systems, the analytical eigenvalue passes through zero as the system goes through a bifurcation, |DEV| for data from continuous systems still approaches 1. This is because DEV is not the analytical eigenvalue of the system, but the dominant eigenvalue of the Jacobian of a discrete impression of the system constructed using lag embedding. DEV shows the same results for at least four types of transitions in continuous systems, further complicating the interpretation of DEV results. Hence, unambiguously identifying the nature of an upcoming transition using DEV is not possible and may warrant further considerations and system-specific knowledge. In complex natural systems, the “ground truth” dynamics are often unknown. DEV will be unable to identify the specific type of transition therein, because it will result in the same trend for multiple types of bifurcations. Thus, the absence of a trend in DEV can probably ensure no transition in the near future, while the presence of trends in DEV can only guide us to dive in further to find out the type of transition.

Preexisting methods to forewarn critical transitions face numerous limitations (Boerlijst et al 2013, Dakos et al 2012a,b, Ditlevsen and Johnsen 2010, Dutta et al 2018, Hastings and Wysham 2010, O’Brien et al 2023). Against this backdrop, DEV has emerged as a new promising EWS that has the potential to predict as well as categorize certain types of bifurcation-induced transitions. The primary strength of DEV lies in the numerical threshold it uses to determine how far a system is from a tipping point. We have shown, using a suite of discrete and continuous models, that DEV possesses certain drawbacks. Thus, one should interpret the DEV results meticulously, taking into account the knowledge of the system in question.

So far, no universal method has been developed to forewarn critical transitions with high accuracy. We support the use of multiple EWSs and other methods in order to predict critical transitions, considering the benefits and limitations of each. DEV is better at classification than spectral EWSs (Bury et al 2020), which are able to differentiate between oscillatory and non-oscillatory bifurcations but fail to make further distinctions. However, even finer classification, that of bifurcations with similar eigenvalue properties, is not possible using DEV. Certain methods, such as machine learning-based EWSs, can distinguish catastrophic bifurcations from non-catastrophic bifurcations and make improved classifications as well (Bury et al 2021, Deb et al 2022, Bury et al 2023). Using multiple strategies in unison may help us not only in weighing the performance of each but also in gaining a multi-pronged understanding of the fate of the system under study.

On the one hand, we observe that DEV has a limited potential to classify transitions. On the other hand, we have also shown that DEV can be extended to provide increasing trends prior to more kinds of bifurcations. Guided by our observations that factors such as the rate of forcing, smoothness of the system, and noise color can influence DEV results, there could exist some natural systems where DEV might not yield expected signals. Regardless, DEV analysis for a system may provide valuable insights about the state and the future trajectory of the system, especially when interpreted using prior knowledge of the system. Future studies can identify the effects of more factors, such as sparsity of data (Kaur and Dutta 2022) on DEV in discrete and continuous systems, and attempt to develop methods that further classify continuous and discontinuous transitions into narrower domains or classes, significantly advancing the research along this direction. The effectiveness of DEV creates new hope for improving our ability to herald and prevent regime shifts, while its limitations assert the need to develop more efficient EWSs and to carefully utilize multiple methods before drawing inferences about the future.

## Declaration of Competing Interest

The authors declare that they have no known competing financial interests or personal relationships that could influence the work reported in this paper.

## Author Contributions

P.S.D. conceived the study. K.K. performed the simulations. All authors analyzed and discussed the results, and wrote the manuscript.

## Funding statement

This work was partially supported by the Science & Engineering Research Board (SERB), Govt. of India [Grant No.: CRG/2022/002788].

## Data availability

Codes and data are available in a GitHub repository (https://github.com/pentathis/DEV-Efficacy).

## Acknowledgments

K.K. acknowledges the Department of Science and Technology, Govt. of India, for the Kishore Vaigyanik Protsahan Yojana (KVPY) fellowship. S.D. acknowledges the Ministry of Education (MoE), Govt. of India, for the Prime Minister’s Research Fellowship (PMRF). P.S.D. acknowledges financial support from the Science & Engineering Research Board (SERB), Govt. of India [Grant No.: CRG/2022/002788].

## SUPPORTING INFORMATION

### S1: Further DEV analysis in multivariate models

In the continuous models containing multiple variables (Table S2, Models 3 and 5), we performed DEV analysis using all the variables and also performed multivariate DEV analysis (*see Main text, Methods*). Figs. S1 and S2 show these results for Hopf bifurcation (Table S2 Model 3) and bifurcation without CSD (Table S2 Model 5), respectively, except for the one variable in each model whose results are given in Fig. 3 (*Main text*). Fig. S3 shows multivariate DEV results for the model with a piecewise smooth Neimark-Sacker bifurcation (Table S3, Model 5).

**Figure S1.**
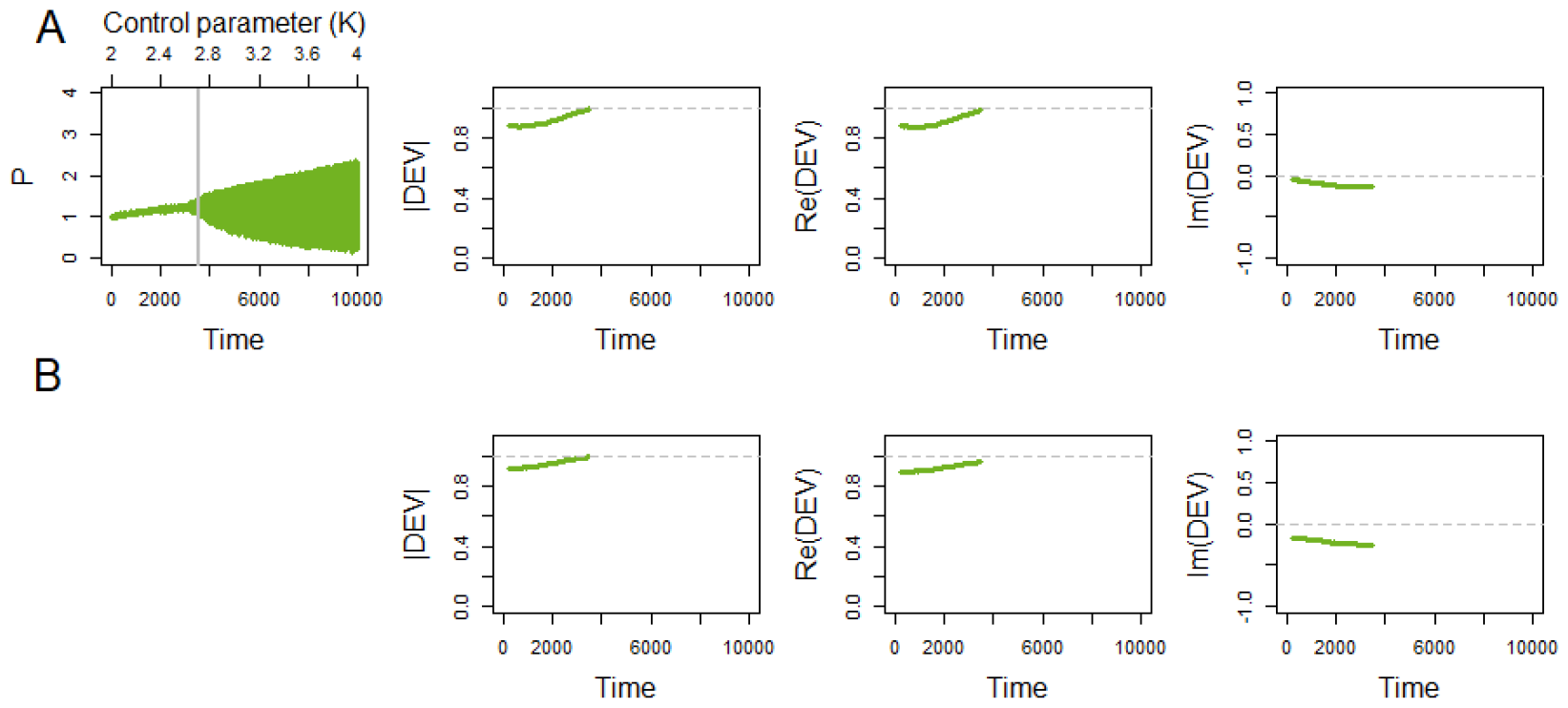
Time series, |DEV|, Re(DEV), and Im(DEV) for Hopf bifurcation (Table S2, Model 3): (A) Univariate DEV for the variable *P*, and (B) Multivariate DEV. The time series is a single realization, whereas the DEV values averaged over 100 realizations. The grey vertical line in the top-left sub-figure marks the transition point. The DEV parameters used are given in Table S6. In both cases, |DEV| → 1, Re(DEV) ↛ 1, and Im(DEV) ↛ 0, successfully predicting the transition.

**Figure S2.**
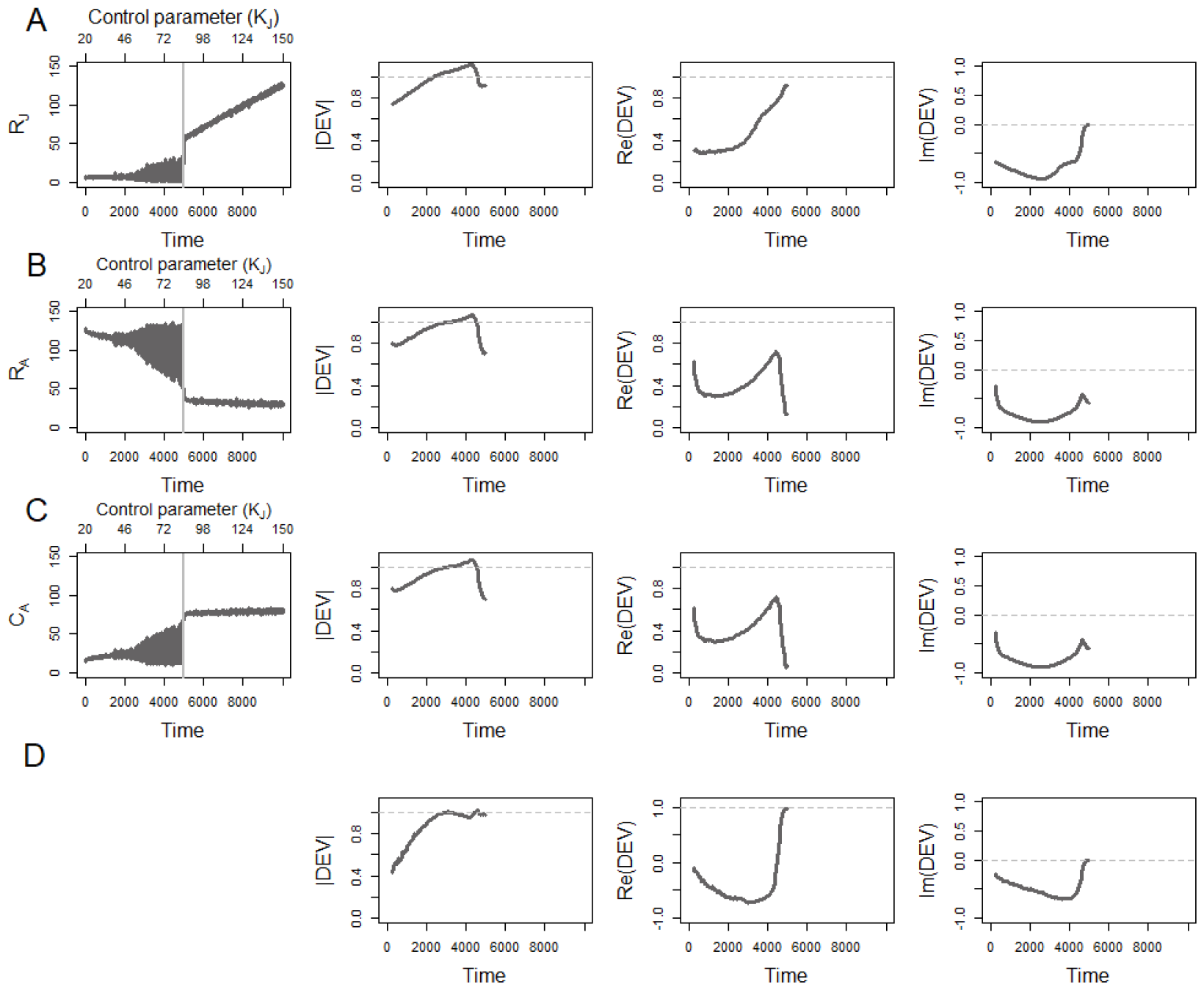
Time series, |DEV|, Re(DEV), and Im(DEV) for bifurcation without CSD (Table S2, Model 5): (A) Univariate DEV for the variable *R*_*J*_, (B) Univariate DEV for the variable *R*_*A*_, (C) Univariate DEV for the variable *C*_*A*_, and (D) Multivariate DEV. The time series is a single realization, whereas the DEV values averaged over 100 realizations. Grey vertical lines in the left-column sub-figures mark the transition points. The parameters used are given in Table S6. As |DEV| reaches one even before the transition and decreases later, the transition is explicitly predicted in none of the cases.

**Figure S3.**
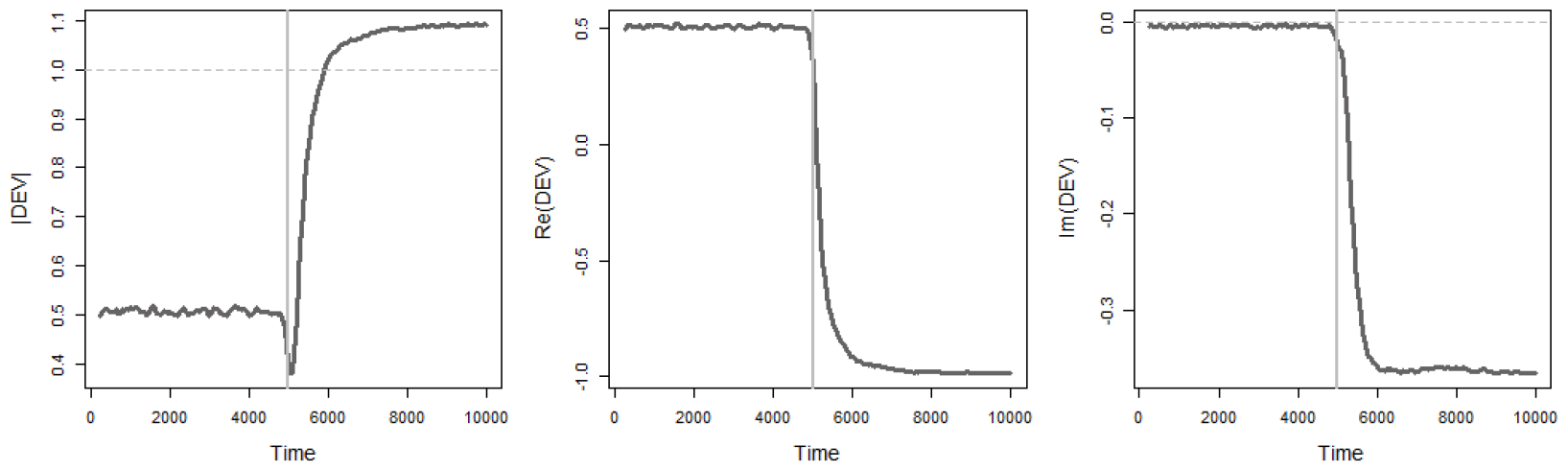
Multivariate |DEV|, Re(DEV), and Im(DEV) for piecewise smooth Neimark-Sacker bifurcation (Table S3, Model 5). The DEV values have been averaged over 100 realizations. Grey vertical lines mark the transition points. The parameters used are given in Table S7. There is no trend in |DEV|, Re(DEV), and Im(DEV) prior to the bifurcation point and a sharp spike at the bifurcation point. Thus, DEV does not predict the transition.

### S2: Sensitivity analysis

Figures S4 and S5 illustrate the sensitivity analysis (see Main text: *Results, Sensitivity analysis*) for the Lotka-Volterra Model (transcritical bifurcation) (Table S1, Model 2) and the reduced Lorenz Model (pitchfork bifurcation) (Table S1, Model 3).

**Figure S4.**
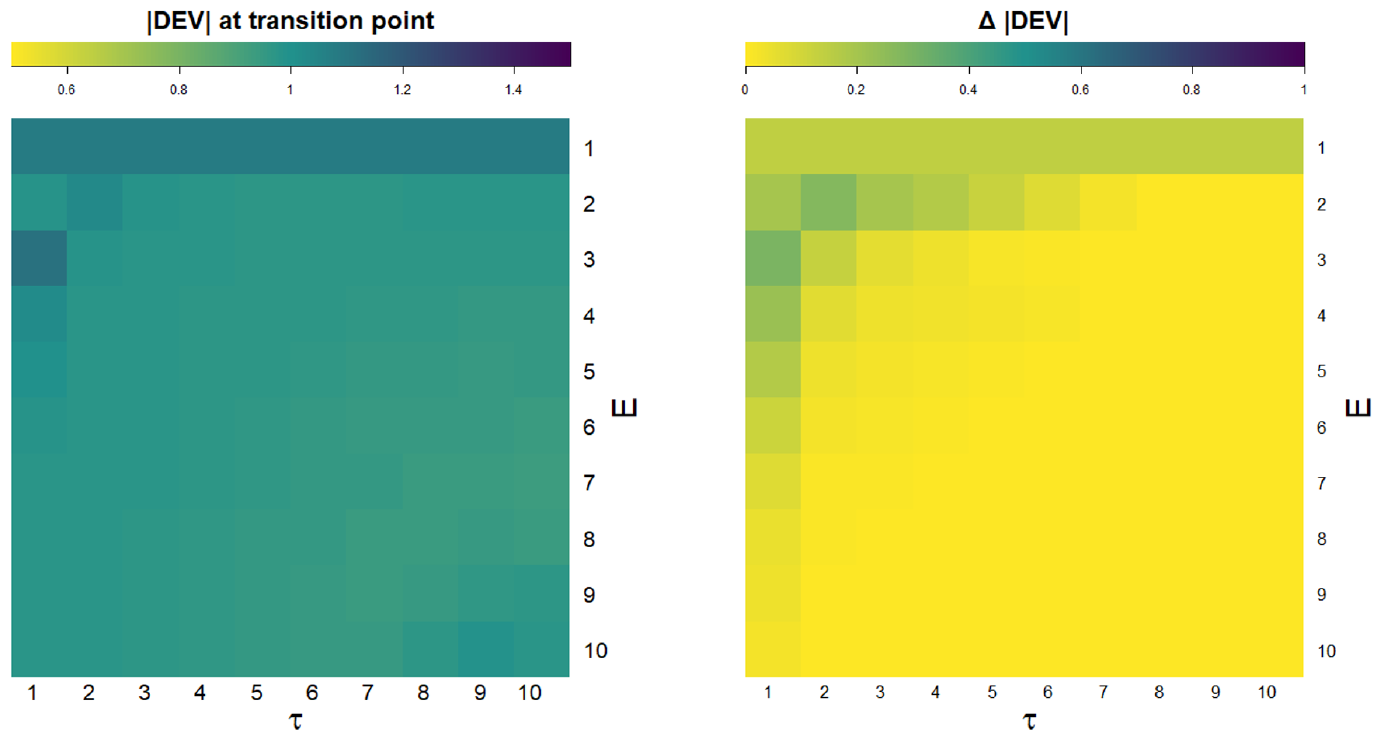
|DEV| at transition point (left) and Δ|DEV| (right) for different combinations of *E* and *τ* for the Lotka-Volterra Model (transcritical bifurcation) (Table S1, Model 2). |DEV| at the transition point remains close to 1 for all values of *E* and *τ*, whereas Δ|DEV| decreases sharply with increasing values of *E* and *τ* .

**Figure S5.**
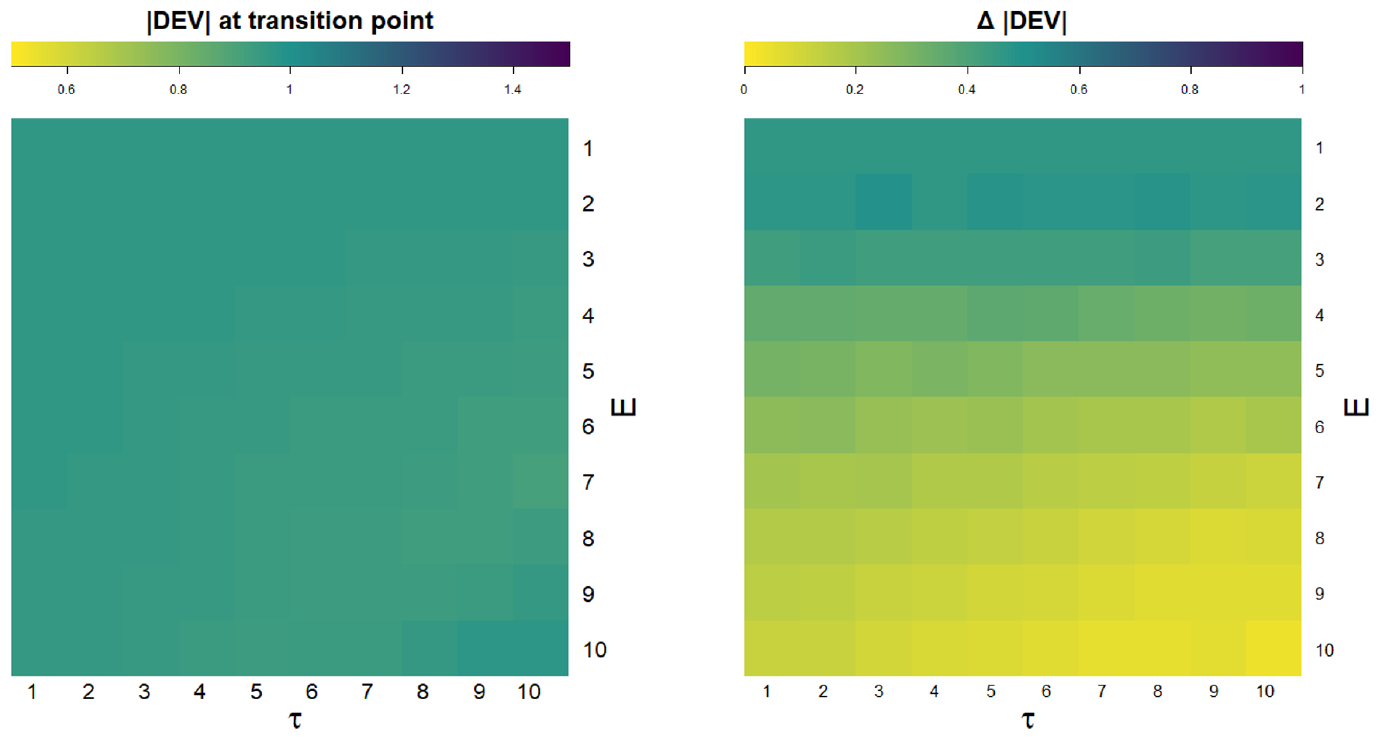
|DEV| at the transition point (left) and Δ |DEV| (right) for different combinations of *E* and *τ* for the reduced Lorenz Model (pitchfork bifurcation) (Table S1, Model 3). |DEV| at the transition point remains close to 1 for all values of *E* and *τ*, whereas Δ |DEV| decreases with increasing values of *E* and decreases at a lesser rate with increasing *τ* .

### S3: Models

Tables S1, S2, S3, and S4 enlist the model equations for discrete models (Noy-Meir 1975, Neubert and Kot 1992, Elabbasy et al 2014, Whitehead and MacDonald 1984), continuous models (Dutta et al 2018), piecewise smooth discrete models (Banerjee et al 2000, De et al 2012) and smooth discrete models (Grziwotz et al 2023) with colored noise, respectively. For each model, the control parameter is given in boldface.

White noise was added to all the variables in the models given in Tables S1, S2 and S3, except in Table S2 Model 5 where it was added to only one variable (*C*_*J*_ ).

Colored noise was added to one variable (the variable for which DEV was evaluated) in each of the models in Table S4, as explained below and in Table S4. When the system entails colored noise, low-frequency noise variations characterize the time series because the next value in the time series is likely to be more similar to the previous one or positively autocorrelated as compared to a time series with white (i.e., uncorrelated) noise (Ripa and Lundberg 1996). Colored noise can be modeled as a first-order autoregressive process. Thus, for every model in Table S4, we consider:

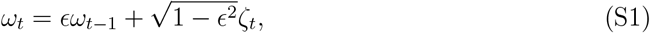

where *ϵ* is the noise correlation and *ζ*_*t*_ is drawn from the standard normal distribution *N* (0, 1). *σ* (Table S4) is a parameter that scales the noise intensity. *ω*_0_ is drawn from the standard normal distribution *N* (0, 1), and 0 ≤ *ϵ <* 1. White noise is the limiting case where *ϵ* = 0.

**Table S1.** Discrete Models

| Deterministic skeleton | Parameters | Values |
| --- | --- | --- |
| 1. Fold bifurcation<br>$N_{t+1} = N_t \exp \left( b - \frac{N_t}{K} \right) - F \frac{N_t^2}{N_t^2 + b^2}$ | $b$<br>$K$<br>**F** | 0.75<br>10<br>**0 to 2** |
| 2. Transcritical bifurcation<br>$x_{t+1} = (r + 1)x_t - rx_t^2 - cx_t y_t$ $y_{t+1} = cx_t y_t$ | $r$<br>**c** | 0.5<br>**1.5 to 0.5** |
| 3. Pitchfork bifurcation<br>$x_{t+1} = (1 + ah)x_t - hx_t y_t$ $y_{t+1} = (1 - h)y_t - hx_t^2$ | $h$<br>**a** | 10<br>**-1 to 1** |

**Table S2.** Continuous models

| Deterministic skeleton | Parameters | Values |
| --- | --- | --- |
| 1. Fold bifurcation |  |  |
| $\frac{dN}{dt} = rN \left(1 - \frac{N}{K}\right) - \frac{cN^2}{b^2 + N^2}$ | $K$-carrying capacity<br>$r$ - maximum growth rate<br>$b$ - half-saturation constant<br>**c - maximum grazing rate** | 10<br>1<br>1<br>**1 – 3** |
| 2. Transcritical bifurcation |  |  |
| $\frac{dN}{dt} = rN \left(1 - \frac{N}{K}\right) - cN$ | $K$ - carrying capacity<br>$r$ - maximum growth rate<br>**c - maximum grazing rate** | 10<br>1<br>**0 – 2** |
| 3. Supercritical Hopf bifurcation |  |  |
| $\frac{dN}{dt} = rN \left(1 - \frac{N}{K}\right) - \frac{aNP}{b + N}$<br>$\frac{dP}{dt} = \frac{eaNP}{b + N} - mP$ | **K - carrying capacity of resource**<br>$r$ - maximum growth rate of resource<br>$a$- maximum grazing rate<br>$b$ - half-saturation constant<br>$e$ - assimilation efficiency of grazer<br>$m$ - mortality rate of grazer | **2 – 4**<br>0.5<br>0.4<br>0.6<br>0.6<br>0.15 |
| 4. Pitchfork bifurcation |  |  |
| $\frac{dN}{dt} = rN(1 - \frac{N}{K})(N - N_c) - cN + I$ | $K$ - carrying capacity<br>**r - maximum growth rate**<br>$c$ - maximum grazing rate<br>$N_c$ - Allee threshold<br>$I$ - immigration rate | 10<br>**0 – 1**<br>0.8<br>5<br>4 |
| 5. Bifurcation without CSD |  |  |
| $\frac{dR_i}{dt} = rR_i(1 - \frac{R_i}{K_i}) - a_iR_iC_i$ ($i = j, A$)<br>$\frac{dC_J}{dt} = ea_AR_AC_A - mC_J - ea_JR_JC_J$<br>$\frac{dC_A}{dt} = ea_JR_JC_J - mC_A$ | $r$ - intrinsic growth rate<br>$m$ - mortality rate<br>$e$ - energy conversion efficiency<br>**K<sub>J</sub> - juvenile carrying capacity**<br>$K_A$ - adult carrying capacity | 1<br>0.1<br>0.4<br>**20 – 150**<br>15 |
| 6. No transition |  |  |
| $\frac{dN}{dt} = rN \left(1 - \frac{N}{K}\right) - \frac{cN^2}{b^2 + N^2}$ | $K$-carrying capacity<br>$r$ - maximum growth rate<br>$b$ - half-saturation constant<br>**c - maximum grazing rate** | 1.9<br>1<br>1<br>**1 – 3** |
| 7. Sharp transition without bifurcation |  |  |
| $\frac{dN}{dt} = rN \left(1 - \frac{N}{K}\right) - \frac{cN^2}{b^2 + N^2}$ | $K$ - carrying capacity<br>$r$ - maximum growth rate<br>$b$ - half-saturation constant<br>**c - maximum grazing rate** | 5.2<br>1<br>1<br>**1 – 3** |

**Table S3.** Discrete models with piecewise smooth bifurcations

| Type of system | Parameters | Values |
| --- | --- | --- |
| Model equations for 1. - 4.: $x_{n+1} = \begin{cases} ax_n + \mu & \text{if } x_n < 0, \\ bx_n + \mu & \text{if } x_n \geq 0. \end{cases}$ | | |
| 1. No bifurcation | a<br>b<br>**$\mu$** | 0.6<br>0.2<br>**-1 to 1** |
| 2. Fold bifurcation | a<br>b<br>**$\mu$** | 1.5<br>0.2<br>**-1 to 1** |
| 3. Period-doubling bifurcation | a<br>b<br>**$\mu$** | -0.5<br>-1.5<br>**1 to -1** |
| 4. Period-1 to chaos | a<br>b<br>**$\mu$** | 0.6<br>-4<br>**-1 to 1** |
| 5. Neimark-Sacker bifurcation | $\begin{bmatrix} x_{n+1} \\ y_{n+1} \end{bmatrix} = \begin{cases} \begin{bmatrix} \tau_L x + y + \mu \\ -\delta_L x \end{bmatrix} & \text{if } x < 0 \\ \begin{bmatrix} \tau_R x + y + \mu \\ -\delta_R x \end{bmatrix} & \text{if } x \geq 0 \end{cases}$ | |
| | $\tau_L$<br>$\tau_R$<br>$\delta_L$<br>$\delta_R$<br>**$\mu$** | -0.1<br>-0.685<br>-0.3<br>1.4<br>**-0.01 to 0.01** |

**Table S4.** Discrete models with colored noise

| Deterministic skeleton + (Noise) | Parameters | Values |
| --- | --- | --- |
| 1. Fold bifurcation<br>$N_{t+1} = N_t \exp \left( b - \frac{N_t}{K} \right) - F \frac{N_t^2}{N_t^2 + b^2} + N\sigma\omega_t$ | $b$<br>$K$<br>**F** | 0.75<br>10<br>**0 to 2** |
| 2. Period-doubling bifurcation<br>$x_{t+1} = 1 - ax_t^2 + y_t + x\sigma\omega_t$ $y_{t+1} = cx_t$ | $c$<br>**a** | 0.3<br>**0.1 to 0.4** |
| 3. Neimarck-Sacker bifurcation<br>$x_{t+1} = (1 + l)x_t - hx_t^2 - \frac{x_t y_t}{1 + bx_t} + x\sigma\omega_t$ $y_{t+1} = -cy_t + \frac{mx_t y_t}{1 + bx_t}$ | $h$<br>$b$<br>$c$<br>$m$<br>**l** | 4<br>0.5<br>2<br>6<br>**3.48 to 3.78** |

### S4: Parameter values used in analyses

Tables S5, S6, and S7 enlist the DEV parameters used for discrete models, continuous models, piecewise smooth discrete models, and discrete models with multiplicative red noise, respectively. The parameters in Table S8 are the same as the ones chosen by Grziwotz et al (2023).

**Table S5.** Parameters chosen for discrete systems

| System | | $w$ | $E$ | $\tau$ |
| --- | --- | --- | --- | --- |
| Fold bifurcation | Table S1 Model 1 | 250 | 2 | 1 |
| Transcritical bifurcation | Table S1 Model 2 | 250 | 2 | 1 |
| Pitchfork bifurcation | Table S1 Model 3 | 250 | 2 | 1 |

**Table S6.** Parameters chosen for continuous systems

| System | | $w$ | $E$ | $\tau$ |
| --- | --- | --- | --- | --- |
| Fold bifurcation | Table S2 Model 1 | 250 | 2 | 1 |
| Transcritical bifurcation | Table S2 Model 2 | 250 | 2 | 1 |
| Hopf bifurcation | Table S2 Model 3 | 250 | 6 | 1 |
| Pitchfork bifurcation | Table S2 Model 4 | 250 | 2 | 1 |
| Bifurcation without CSD | Table S2 Model 5 | 250 | 6 | 1 |
| No transition | Table S2 Model 6 | 250 | 2 | 1 |
| Transition without bifurcation | Table S2 Model 7 | 250 | 2 | 1 |

**Table S7.** Parameters chosen for piecewise smooth discrete systems

| System | | $w$ | $E$ | $\tau$ |
| --- | --- | --- | --- | --- |
| No bifurcation | Table S3 Model 1 | 250 | 2 | 1 |
| Fold bifurcation | Table S3 Model 2 | 250 | 2 | 1 |
| Period-doubling bifurcation | Table S3 Model 3 | 300 | 2 | 1 |
| Period-1 to chaos | Table S3 Model 4 | 250 | 2 | 1 |
| Neimark-Sacker bifurcation | Table S3 Model 5 | 250 | 4 | 1 |

**Table S8.** Parameters chosen for discrete systems perturbed with colored noise

| System | | $w$ | $E$ | $\tau$ |
| --- | --- | --- | --- | --- |
| Fold bifurcation | Table S4 Model 1 | 250 | 2 | 1 |
| Period bifurcation | Table S4 Model 2 | 100 | 3 | 1 |
| Neimark-Sacker bifurcation | Table S4 Model 3 | 100 | 6 | 1 |

## Notes

https://github.com/pentathis/DEV-Efficacy

### Competing Interest Statement

The authors have declared no competing interest.

